# Comparison of SARS-CoV-2 entry inhibitors based on ACE2 receptor or engineered Spike-binding peptides

**DOI:** 10.1101/2023.01.05.522964

**Authors:** George N. Llewellyn, Hsu-Yu Chen, Geoffrey L. Rogers, Xiaoli Huang, Philip J. Sell, Jill E. Henley, Paula M. Cannon

**Affiliations:** Department of Molecular Microbiology and Immunology, Keck School of Medicine of the University of Southern California, Los Angeles, CA; The Hastings Foundation and The Wright Foundation Laboratories, Keck School of Medicine of the University of Southern California, Los Angeles, CA

## Abstract

With increasing resistance of SARS-CoV-2 variants to antibodies, there is interest in developing entry inhibitors that target essential receptor binding regions of the viral Spike protein and thereby present a high bar for viral resistance. Such inhibitors can be derivatives of the viral receptor, ACE2, or peptides engineered to interact specifically with the receptor-binding pocket. We compared the efficacy of a series of both types of entry inhibitors, constructed as fusions to an antibody Fc domain. Such a design can increase protein stability and act to both neutralize free virus and recruit effector functions to clear infected cells. We tested the reagents against prototype variants of SARS-CoV-2, using both Spike pseudotyped VSV vectors and viral plaque assays. These analyses revealed that an optimized ACE2 derivative could neutralize all variants we tested with high efficacy. In contrast, the Spike-binding peptides had varying activities against different variants, with resistance observed for the Spike proteins from Beta, Gamma and Omicron. The resistance mapped to mutations at Spike residues K417 and N501 and could be overcome for one of the peptides by linking two copies in tandem, effectively creating a tetrameric reagent in the Fc fusion. Finally, both the optimized ACE2 and tetrameric peptide inhibitors provided some protection to human ACE2 transgenic mice challenged with the SARS-CoV-2 Delta variant, which typically causes death in this model within 7-9 days.

**Importance:** The increasing resistance of SARS-CoV-2 variants to therapeutic antibodies has highlighted the need for new treatment options, especially in individuals who do not respond to vaccination. Receptor decoys that block viral entry are an attractive approach because of the presumed high bar to developing viral resistance. Here, we compare two entry inhibitors based on derivatives of the ACE2 receptor or engineered peptides that bind to the receptor binding pocket of the SARS-CoV-2 Spike protein. In each case, the inhibitors were fused to immunoglobulin Fc domains, which can further enhance therapeutic properties, and compared for activity against different SARS-CoV-2 variants. Potent inhibition against multiple SARS-CoV-2 variants was demonstrated *in vitro*, and even relatively low single doses of optimized reagents provided some protection in mouse models, confirming their potential as an alternative to antibody therapies.

## Introduction

SARS-CoV-2, the virus that causes COVID-19, has spread globally since first identified in Wuhan, China in 2019 [1]. By Fall 2022, more than 637 million cases and 6.62 million deaths have been reported [2]. Contributing to this pandemic has been the emergence of distinct viral variants, of which the World Health Organization has designated 5 as variants of concern: Alpha (B.1.1.7, first detected in the UK in 2020), Beta (B.1.351, from South Africa in 2020), Gamma (P.1, from Brazil in 2021), Delta (B.1.617.2, from India in 2021) and Omicron (B.1.1.539, from multiple countries in 2021). Most recently, Omicron BA.1 has given rise to several subvariants such as BA.2, BA.4 and BA.5, which have contributed to new waves of infections [3].

To combat this public health emergency, vaccines, monoclonal antibody treatments and antiviral drugs have been rapidly developed [4, 5]. While vaccines have proven successful at preventing serious illness and death, they do not absolutely prevent infection. In particular, early studies reported a diminished ability of vaccine-induced antibodies to neutralize the Beta and Gamma variants [6, 7] and recent studies have shown similar escape from vaccine-elicited antibodies by Omicron subvariants [8]. More recently, viruses in the Omicron lineage have been shown to be capable of causing breakthrough infections in both previously vaccinated or infected individuals [9], and the extensive mutations present in the Omicron Spike protein have prevented the effectiveness of antibody therapies [10–12]. This underscores the need for alternate, broadly-acting vaccines and therapeutics with activity against both current and future viral variants [13].

SARS-CoV-2 enters cells following the interaction of its Spike protein with cell surface angiotensin-converting enzyme 2 (ACE2) [1]. Viral receptors can be repurposed as entry inhibitors, being expressed both as soluble receptor domains or further stabilized by fusion to other proteins such as immunoglobulin (Ig) Fc domains [14,15]. The underlying assumption is that a virus would not be able to evolve resistance to an inhibitor that mimics the interaction with its cellular receptor without a significant loss of fitness. Both soluble ACE2 and ACE2-Ig fusions have been reported as effective inhibitors of entry directed by the Spike proteins from SARS-CoV-1 [16–19] and SARS-CoV-2 [16, 17, 20–32]. To prevent interference with the normal function of ACE2 in regulating blood pressure, these antiviral reagents also usually contain mutations in the protein’s catalytic domain [18]. Studies have also identified additional mutations in ACE2 that enhance binding to the SARS-CoV-2 Spike, increasing neutralization in *in vitro* assays by 100 to 300-fold [20–22, 28, 33, 34]. Of note, binding-enhanced ACE2-Ig constructs have shown activity against different variants of concern, including Omicron [26, 30, 34–37]. Finally, ACE2-Ig constructs have been reported to provide some protection following SARS-CoV-2 challenge of hamsters [20, 30] and human ACE2 transgenic mice [22, 24, 30, 32, 34, 35, 37].

An alternative approach to inhibiting SARS-CoV-2 entry has been to use small Spike-binding peptides (SBP), engineered to target the essential receptor binding domain (RBD) of the protein [38–40]. These SBPs can prevent infection at picomolar levels *in vitro*, and are protective when given to mice as either the peptides alone [39], when fused to an Fc domain [39], or when trimerized to match the oligomeric form of the Spike protein [40].

In the present study, we compared the potency and breadth of activity of a binding-enhanced ACE2-Ig inhibitor and a series of constructs comprising SBPs fused to antibody Fc domains (SBP-Ig). We tested the reagents against pseudoviruses containing SARS-CoV-2 Spike proteins from both the early emerging D614G variant [41] and representatives of each of the major variants of concern. We found that while the enhanced ACE2-Ig derivative could block entry by all Spike proteins we tested with similar picomolar efficacy, the two SBP-Ig inhibitors were sensitive to the mutations acquired by the Beta, Gamma, and Omicron variants. Interestingly, presentation of one of the small peptides as a tandem repeat was able to overcome this resistance, restoring broad activity to this class of reagents, and agreeing with previous reports of the superior activity of multimeric peptide designs [40]. Finally, single injections of both the modified ACE2-Ig construct and a tandem SBP-Ig reagent showed some protection in mice challenged with the Delta variant, with the multimerized SBP-Ig giving a higher survival rate.

## Results

### Optimization of ACE2-Ig for neutralization of the SARS-CoV-2 Spike protein

To identify an optimal ACE2-Ig entry inhibitor, we first compared constructs comprising residues 1-740 of the ACE2 ectodomain fused to IgG1 Fc and a shorter version, truncated at residue 615, that excluded the ACE2 collectrin-like domain. In both cases, a catalytically dead ACE2 variant was used that does not cleave the angiotensin-(1-8) peptide [18]. Previously, an ACE2-Ig construct comprising residues 18 to 740 was found to have superior neutralizing activity than a 18-615 version against pseudoviruses carrying the original SARS-CoV-2 Wuhan Spike protein [17]. We also observed that our longer 1-740 version was superior to the 1-615 version in neutralization studies using pseudoviruses carrying the D614G Spike protein, which was an early dominant variant of SARS-CoV-2, with enhanced infectivity and spread [42, 43] (**Supplementary Figure 1**).

The potency of ACE2 receptor decoys can be increased by mutations that enhance binding to the RBD of the SARS-CoV-2 Spike [20–22, 28, 33–35]. We selected the V2.4 mutation group (T27Y, L79T and N330Y), which was reported to enhance binding by 35-fold, and protected mice from an infectious challenge when delivered as either the soluble protein or fused to an Fc domain [28, 35]. In addition, we attempted to further enhance the ACE2-Ig constructs by inclusion of a heptad repeat-2 (HR2) peptide fusion inhibitor [44] at the C-terminus of the Fc domain (**Fig. 1A**).

**Figure 1.**
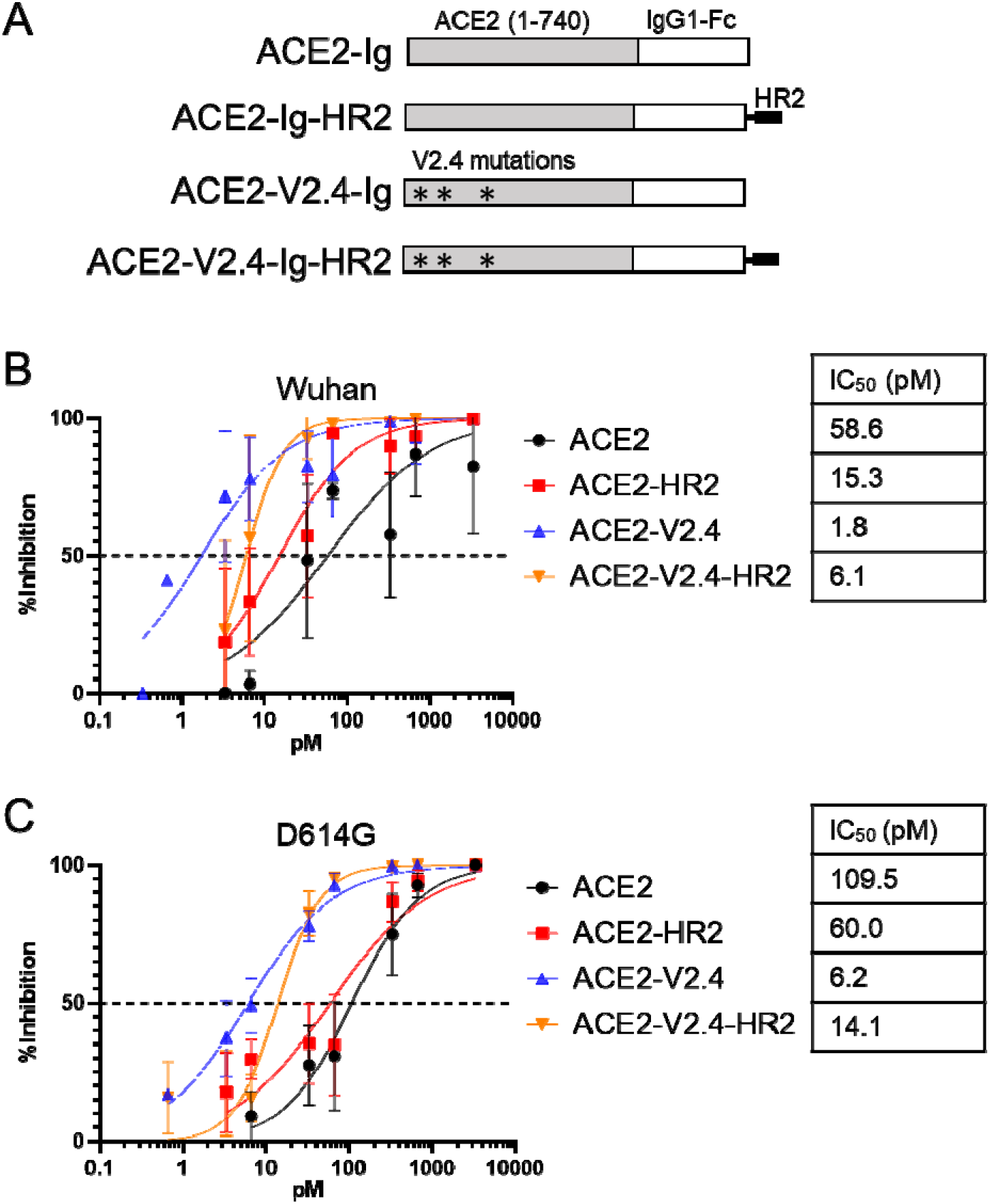
ACE2 binding enhancing mutations, but not HR2, increase neutralization of SARS-CoV-2 Spike. **A)** Schematic of different ACE2-Ig constructs. **B)** Neutralization of VSV vectors pseudotyped with the Wuhan variant Spike protein by indicated amounts of ACE2-Ig constructs. Error bars are SEM from n=3 independent experiments and mean IC_50_ values are indicated. **C)** Neutralization assay for VSV pseudovirus containing the D614G Spike variant. Error bars are SEM from n=4 independent experiments and mean IC_50_ values are indicated.

To test these various entry inhibitors, we used a Spike pseudovirus system that we have previously optimized [45]. Here, SARS-CoV-2 Spike proteins are incorporated into replication-incompetent VSV vectors that express luciferase upon entry, and the vectors are used to transduce HeLa cells expressing ACE2. Using this system, we compared the ability of the series of ACE2-Ig constructs to inhibit entry directed by pseudoviruses containing the Spike proteins from either the original Wuhan isolate or the D614G variant (**Fig. 1B-C**). We found that the V2.4 mutations enhanced inhibition 18 to 33-fold against the D614G and Wuhan pseudoviruses, respectively. In contrast, the HR2 peptide resulted in only a small enhancement for the non-modified ACE2-Ig construct, and slightly decreased inhibition by ACE2-V2.4-Ig. As a result, we moved forward with ACE2-V2.4-Ig as an optimal design.

### Spike-binding peptides fused to Fc domains are potent entry inhibitors

A series of computationally designed peptides have been described that bind to different regions of the surface of the Spike RBD surrounding the ACE2 binding site, and which inhibit SARS-CoV-2 infection of Vero E6 cells [38]. From these, we selected peptides LCB1 and LCB3, which were especially potent and inhibited infection by the Wuhan variant at picomolar levels. We fused each peptide to an IgG1 Fc domain to create LCB1-Ig and LCB3-Ig fusion constructs which, for simplicity, we re-name here as Spike binding peptide 1 (SBP1)-Ig and SBP2-Ig. A similar LCB1-Fc construct has previously been described [39], which protected human ACE2 transgenic mice from infection by the Wuhan strain of SARS-CoV-2 if given as an IP injection of 250μg, either 1 day pre- or 1 day post-exposure. We compared the efficiency of each of the two SBP-Ig constructs against both the original and binding-enhanced ACE2-Ig constructs, using D614G Spike pseudovirus neutralization assays. Here, we observed that both SBP-Ig constructs had similar potency as the enhanced ACE2-V2.4-Ig construct, in the 11-14 picomolar range (**Fig. 2**).

**Figure 2.**
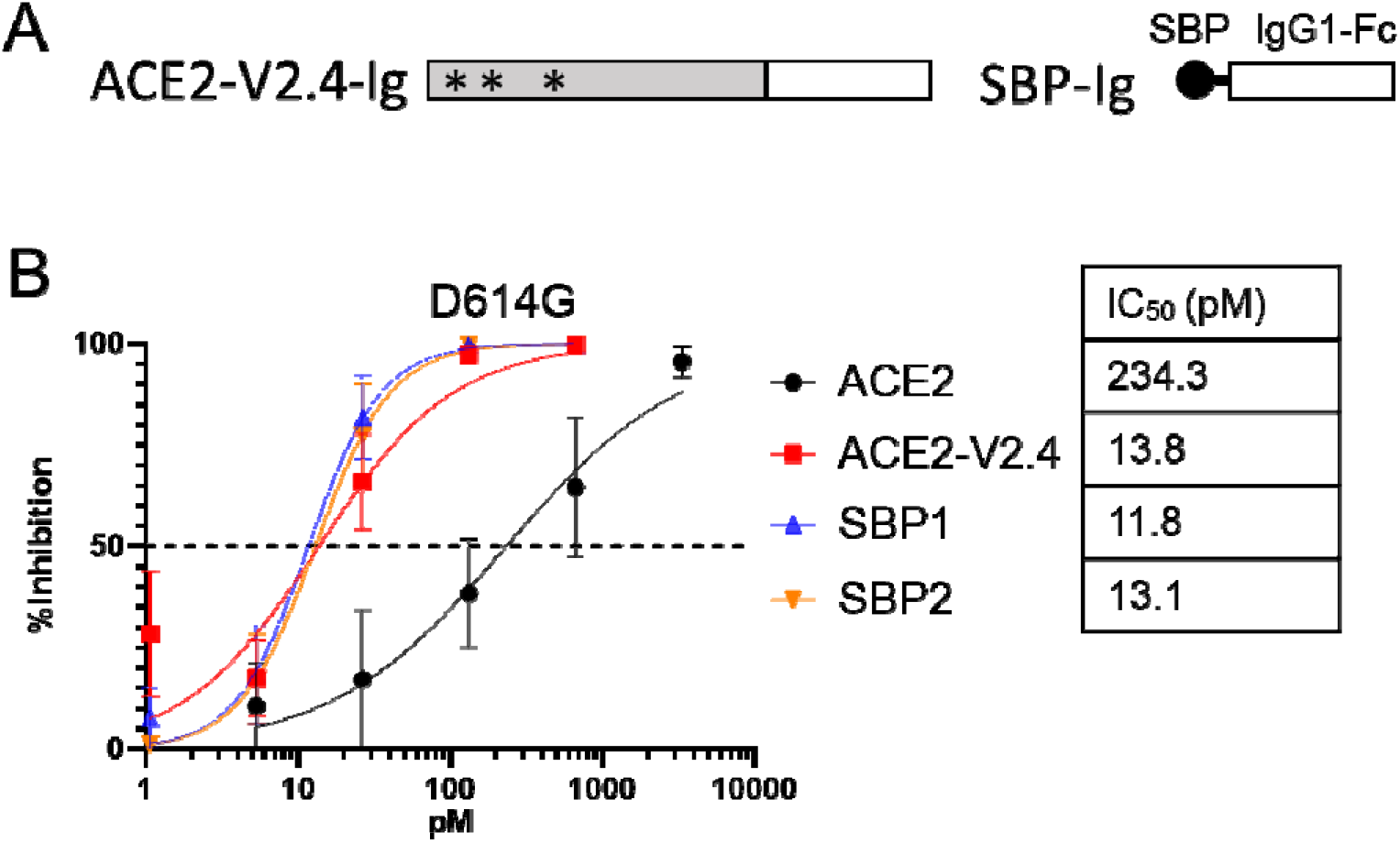
Spike-binding peptides attached to IgG1 Fc domain inhibit D614G pseudovirus. **A)** Schematic of SBP-Ig and ACE2-V2.4-Ig constructs. **B)** Neutralization assay for VSV pseudovirus containing the D614G Spike variant. Error bars are SEM from n=4 independent experiments and mean IC_50_ values are indicated.

### Neutralization of Spike proteins from major SARS-CoV-2 variants

We next compared the panel of ACE2-Ig and SBP-Ig-based entry inhibitors against pseudoviruses containing Spike proteins from 6 major SARS-CoV-2 variants: Alpha, Beta, Epsilon, Gamma, Delta and Omicron (**Fig. 3**). We found that ACE2-V2.4-Ig potently inhibited all 6 variants, with an IC_50_ range of 6.5 to 33.5 pM and was always more effective than the unmodified ACE2-Ig construct. In contrast, the two SBP-Ig constructs showed differences in their ability to neutralize Spike variants. Although both inhibited the Alpha, Epsilon and Delta Spikes, with SBP1-Ig especially being the most potent of all the constructs we tested against these variants, neither SBP-Ig had activity against the Beta, Gamma and Omicron Spikes at the tested concentrations (**Fig. 3**). This is consistent with observations that RBDs containing mutations characteristic of the Beta, Gamma and Omicron Spikes (K417N, E484K and N501Y) increased dissociation rates of the LCB1 peptide bound to the RBD by over 100-fold, [40].

**Figure 3.**
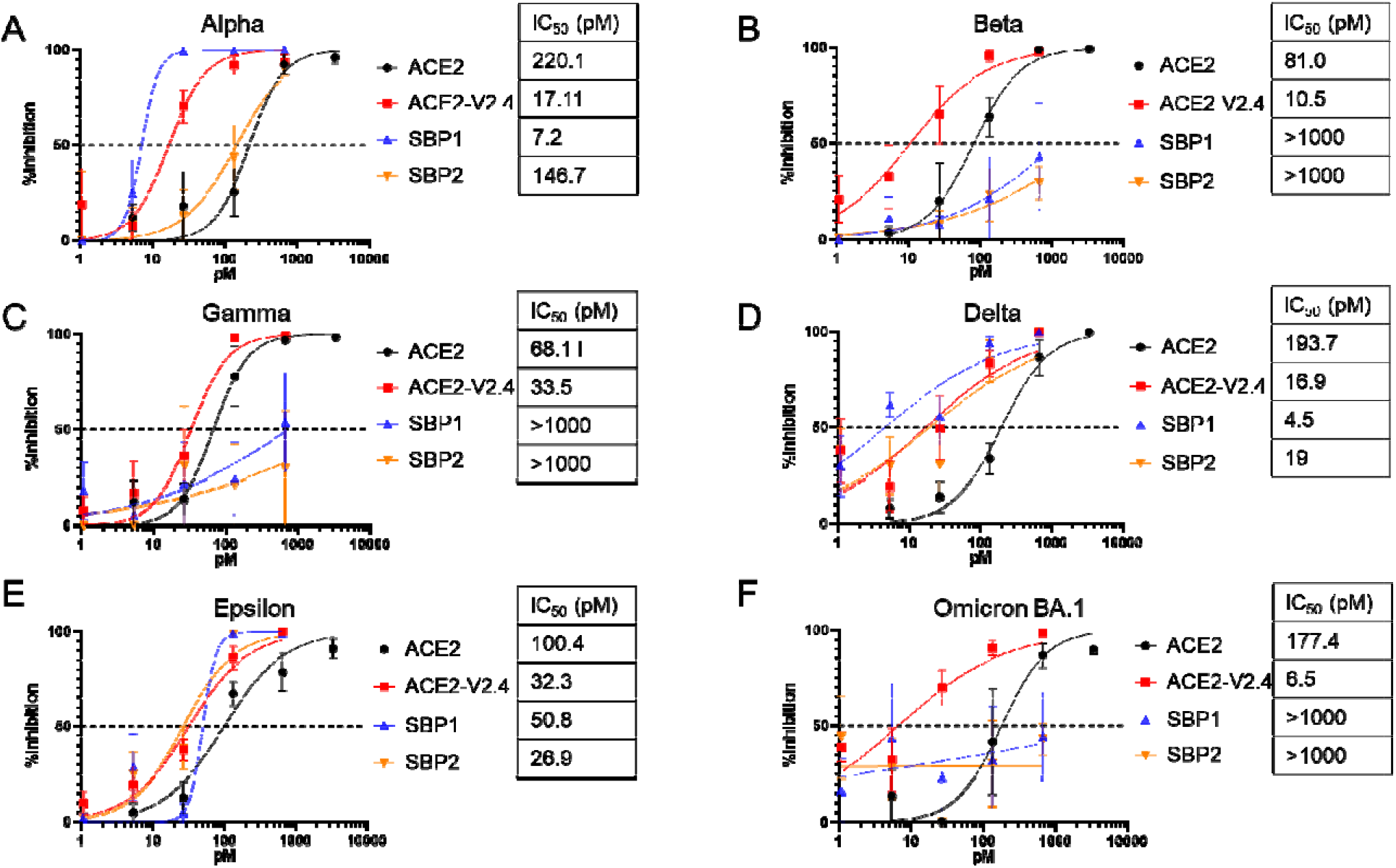
Inhibition of major SARS-CoV-2 variants. Neutralization assays for VSV pseudoviruses containing Spike proteins from SARS-CoV-2 variants Alpha **(A)**, Beta **(B),** Gamma **(C)**, Delta **(D)**, Epsilon **(E)** and Omicron BA.1 **(F).** Error bars are SEM from n=4 independent experiments and mean IC_50_ values are indicated.

### Effect of entry inhibitors on replication of selected SARS-CoV-2 variants

We assessed whether the entry inhibitors maintained their activity against replication competent SARS-CoV-2. Using a qualitative plaque assay, we examined the efficacy of ACE2-Ig, ACE2-V2.4 and SBP2-Ig against a subset of viral variants, informed by the pseudovirus assay observations i.e. Wuhan, Beta and Delta. Similar to the pseudovirus assays, we found that ACE2-Ig and ACE2-V2.4-Ig inhibited all 3 variants, while SBP2-Ig inhibited Wuhan and Delta, but not the Beta variant (**Supplemental Figure 2**). These results therefore confirm the usefulness of the pseudovirus system as a model to test SARS-CoV-2 entry inhibitors.

### Spike protein residues K417 and N501 contribute to SBP-Ig resistance

The lack of ability of the two SBP-Ig constructs to inhibit the Beta, Gamma or Omicron pseudoviruses at the concentrations tested corresponds with the ability of these variants to also resist neutralization by antibodies in vaccinated individuals [6–8]. Sequence alignments of RBDs reveals mutations that are conserved within the resistant variants but are not present in the susceptible variants we tested, specifically residues 417, 484 and 501 (**Supplemental Figure 3**). These residues have also been implicated in monoclonal antibody escape [7]. In addition, crystal structures of the two SBPs complexed with the Wuhan Spike protein [38], show that residues K417, E484 and N501 are within the SBP binding pocket (**Supplemental Figure 4**).

We therefore introduced mutations at these residues into the D614G variant and analyzed the effect on Spike pseudovirus neutralization by the SBP-Ig constructs. This revealed that the K417N or K417T substitutions alone were sufficient to confer resistance against SBP2-Ig. In contrast, resistance to SBP1-Ig required the double substitution of K417N and N501Y, with neither single mutation alone being sufficient (**Figure 4**). This agrees with previous pseudovirus studies using a modified truncated LCB-1 peptide, which implicated both mutations K417N and N501Y in Spike resistance [46].

**Figure 4.**
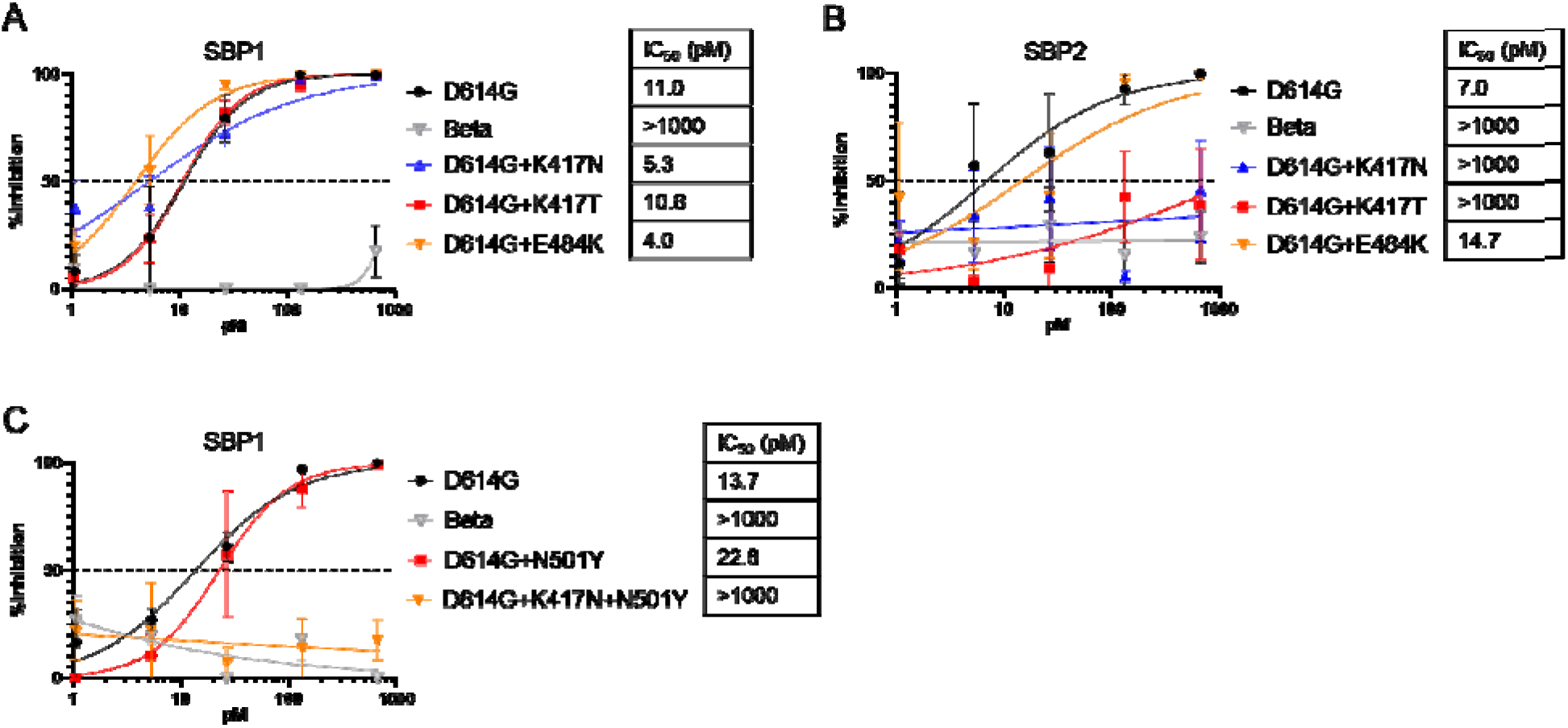
Importance of K417N/T and N501Y mutations in SBP-Ig resistance. Neutralization assays for VSV pseudoviruses containing indicated Spike proteins. D614G was used as a positive control and Beta variant B.1.351 as a negative control. Single point mutations K417N, K417T or E484K were introduced into D614G Spike and tested for sensitivity to (**A)** SBP1-Ig, and (**B)** SBP2-Ig. **(C)** Point mutations N501Y and K417N+N501Y were introduced into D614G Spike and tested against SBP1-Ig. Error bars are SEM from n=3 independent experiments and mean IC_50_ values are indicated

### Tandem repeats of SBP-1 overcome resistance by the Beta, Gamma and Omicron Spike proteins

Multiplexing these SBPs as linked trimeric peptide chains, or by fusing the peptides to trimerization domains, was reported to enhance both Spike-binding affinity and avidity [40]. In addition, some multimers overcame the resistance exhibited by Beta and Gamma variants against single copies of the peptides [40]. We therefore examined the function of multiplexed SBPs in the context of the Fc fusion design, creating tandem duplications of the individual SBP1 and SBP2 reagents, as well as mixed combinations (**Fig. 5A**). Because Fc domains naturally dimerize, these constructs are expected to present tetramers of the peptides. In an initial test against the sensitive D614G and Delta variants, we observed minimal effects of adding a second peptide (**Supplemental Figure 5**), indicating that these designs are well-tolerated. For example, both SBP1-Ig and its duplicated version, SBP1-1-Ig, had similar IC_50_ values and remained the most potent constructs against these variants.

**Figure 5.**
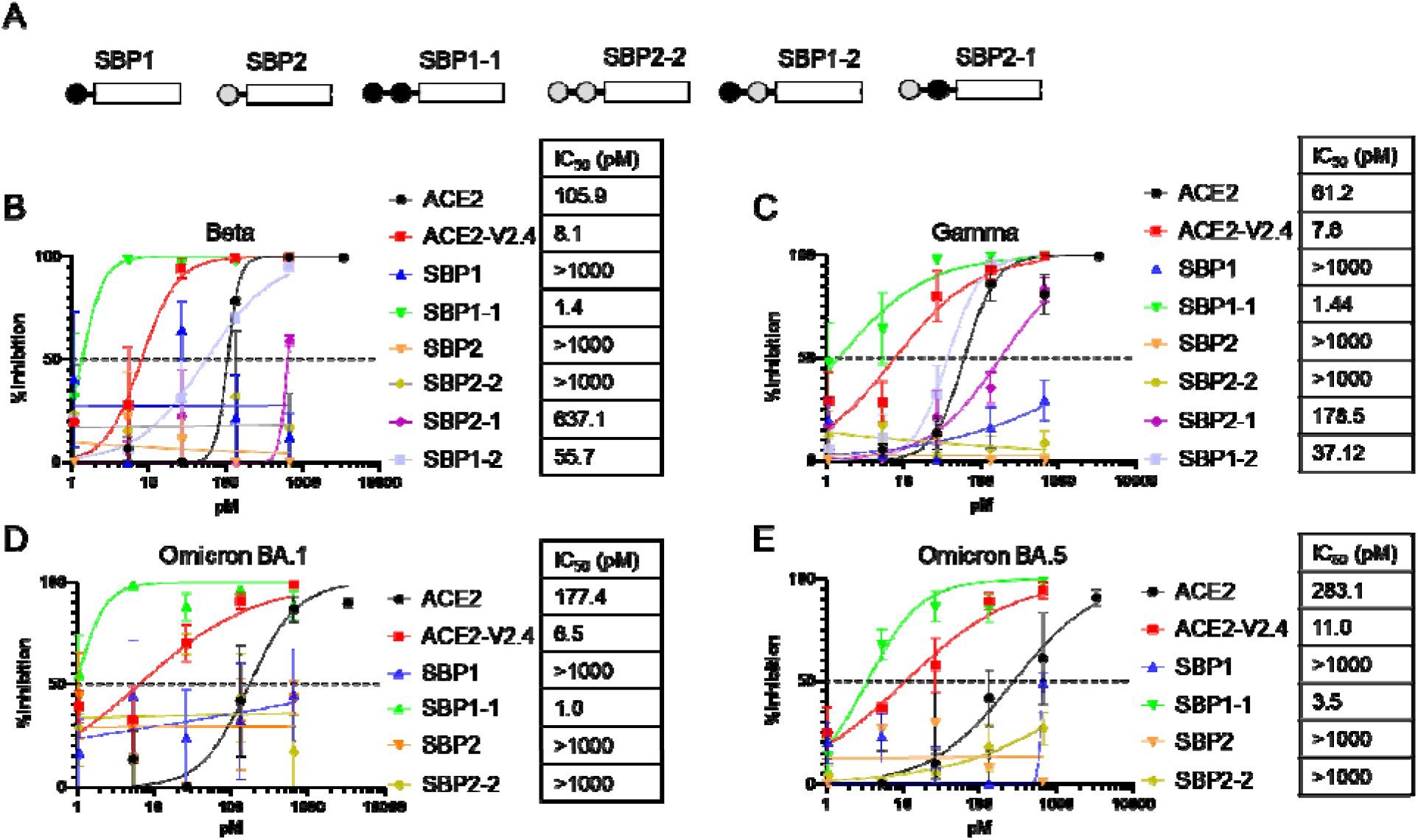
Effects of tandem combinations of SBP1 and SBP2 on neutralization of different Spike variants. **(A)** Schematic of various SBP-Ig constructs. Neutralization assays with indicated SBP-Ig inhibitors were performed against Spike pseudoviruses for **(B)** Beta, **(C)** Gamma, **(D)** Omicron BA.1 and **(E)** Omicron BA.5. Error bars are SEM from n=3 independent experiments and mean IC_50_ values are indicated

We tested the tandem repeat constructs against the SBP-resistant variants Beta, Gamma and the Omicron subvariants BA.1 and BA.5 (**Fig. 5**). Here, we found that a duplication of SBP1, but not SBP2, could overcome the resistance in these Spike proteins. Moreover, SBP1-1-Ig was observed to be the most potent construct tested, having activity in the range of 1.0-3.5 pM, and surpassing the neutralization activity of ACE2-V2.4-Ig. Finally, for the mixed SBP1 and 2 combinations, we also observed increased activity against the Beta and Gamma Spikes, with SBP1 at the N-terminus being the more potent design. These results therefore demonstrate the ability of multiplexed peptide designs in the context of Fc fusions to enhance the potency of peptide-based inhibitors, suggesting a way to limit future evolution of viral resistance.

### Evaluation of entry inhibitors in Delta variant challenge of ACE2 transgenic mice

The most potent and broad inhibitors we identified in our *in vitro* neutralization studies were ACE2-V2.4-Ig and SBP1-1-Ig. We therefore next compared their ability to protect human ACE2 transgenic mice [47] from the health consequences of SARS-CoV-2 infection. As a rigorous test, we selected to use the Delta variant, since this causes rapid weight loss and death in infected animals. The Omicron variant, which in 2022 was the major variant in circulation, was found to be much less pathogenic in this model (data not shown), in agreement with other reports [48].

We first evaluated ACE2-V2.4-Ig. Mice were inoculated intranasally with 1 x 10^3^ PFU of the Delta variant, followed with a single intraperitoneal injection of 10μg of ACE2-V2.4-Ig one day later. In some mice, a second dose was also given at day 5. Mice were observed daily and euthanized if showing significant health deterioration, including losing more than 20% of body weight (**Supplementary Figure 6**). For the single dose group, we observed some protection, with 38% of mice surviving compared to only 5% in the control group (**Fig. 6A**). The group receiving a second dose showed a higher survival rate of 60%.

**Figure 6.**
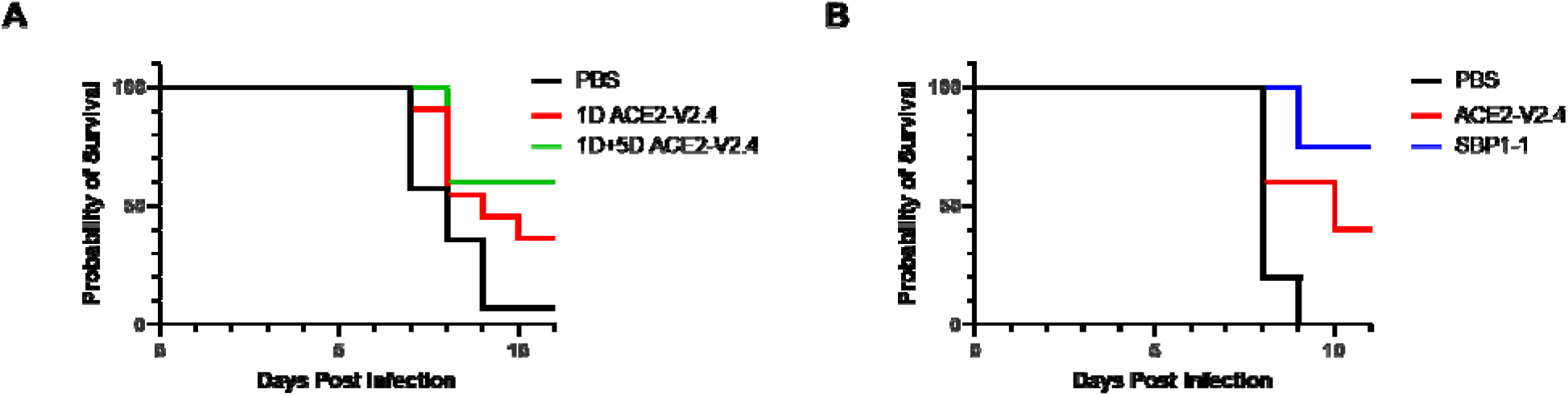
Ability of ACE2-V2.4-Ig and SBP1-1-Ig to protect ACE2 mice from SARS-CoV-2 Delta challenge. Mice were infected intranasally with 10^3^ PFU of Delta variant and monitored daily for weight. Mice were euthanized if more than 20% body weight was lost, or health deteriorated. (**A)** Mice were injected IP with 10μg ACE2-V2.4-Ig, at either 1 day (n=16) or 1 plus 5 days (n=5) post infection or were injected with PBS controls (n=14). (**B)** Mice were injected IP at 12 hours post-infection with 20μg ACE2-V2.4-Ig (n=5) or the molar equivalent for SBP1-1-Ig (7.9μg) (n=4), or a PBS control (n=5).

We then compared ACE2-V2.4-Ig to the tandem repeat construct SPB1-1-Ig, which had the most potent activity against all of the variants tested *in vitro*, including Delta (**Supplementary Figure 5C**). For this experiment, mice were given either a single dose of 20μg of ACE2-V2.4-Ig or the molar equivalent dose of SBP1-1-Ig (7.9μg), 12 hours post infection. In this regimen, ACE2-V2.4-Ig allowed 40% of the mice to survive, while SBP1-1 was more effective, with 75% of the mice surviving after the single dose (**Fig. 6B**).

## Discussion

There is an urgent need for both therapeutic and preventative treatments for COVID-19 to protect against both current and future variants of SARS-CoV-2 that could evolve. This need is illustrated especially by recent members of the Omicron lineage that are resistant to all current monoclonal therapies [10–12]. Towards that goal, entry inhibitors based on, or mimicking, how the ACE2 receptor is recognized by the viral Spike protein are expected to provide a large barrier to viral resistance. Indeed, ACE2-Ig has already been shown to be more resistant to the mutations found in Alpha, Beta, Gamma, Delta and Epsilon variants when compared to monoclonal antibodies [49].

In this study, we directly compared two alternative approaches to entry inhibitors. We used a modified ACE2 domain comprising residues 1-740 and mutations (V2.4) that enhance binding to the SARS-CoV-2 Spike [28], and two potent engineered small Spike-binding peptides [12]. We displayed each inhibitor as a fusion to the IgG1 Fc domain, in a strategy that creates bivalency and is expected to both enhance protein stability and support antibody effector functions that could also clear infected cells. Using a neutralization assay based on Spike VSV pseudoviruses, we found that each of these inhibitors was active at low picomolar concentrations. ACE2-V2.4-Ig proved to be the most broadly inhibitory, neutralizing all SARS-CoV-2 variants that we tested with similar efficacy (6.5-33pM). In contrast, the two SBP-Ig constructs had a narrower range, being susceptible to mutations found in the Spike proteins from Beta, Gamma and Omicron, and specifically K417N and N501Y. However, this resistance could be overcome using a tandem combination of two SBP peptides, where at least the N-terminus peptide was SBP1.

Notably, the most effective antibody therapies have IC_50_ values in the range of 6-7pM against most variant pseudoviruses [12]. In comparison, we found that ACE2-V2.4-Ig was effective in such assays in a similar range (6.5-33pM) against all major variants. The multiplexed SBP1-1-Ig construct was even more potent in pseudovirus assays, neutralizing in the range of 1.0-3.5pM and being effective against even the Omicron BA.5 variant (3.5pM), for which no current antibodies work. This shows that both inhibitor designs hold promise as potent and broad virus neutralizing reagents.

We also compared the abilities of ACE2-V2.4-Ig and SBP1-1-Ig to protect human ACE2 transgenic mice against challenge with the SARS-CoV-2 Delta variant. In this model, we found Delta to be aggressive, causing severe weight loss or death in untreated animals by 7-9 days. To evaluate each construct, we used lower doses than typically given in similar studies, but still found that ACE2-V2.4-Ig protected about 40% of mice from death following a single IP dose of 10-20μg given 1 day post infection. For SBP1-1-Ig, 75% of mice were protected following a single intranasal dose of 7.9μg (the molar equivalent of 20μg of ACE2-V2.4-Ig), also given one day post infection.

Our mouse challenge studies agree with and extend previous studies using ACE2 derivatives, although we achieved protection using much lower doses. For example, a study using an alternate ACE2-Ig, containing different binding enhancing mutations (MDR504 hACE2-Fc), was given to Ad5-ACE2 transduced mice at a dose of 360μg, 4 hours before challenge with the original Wuhan virus, then followed for 3 days before necropsy, or for 7 days to monitor survival. [22]. The authors saw a decrease in viral load and complete survival through the 7 days of the study, although this was a short timeframe to measure survival, since even untreated animals had almost 50% survival at that point. Another study gave mice daily injections of ACE2-V2.4-Ig, at a dose of 10-15 mg/kg per day (roughly 200-300μg/day) for up to 14 days post infection. This study reported 50% survival for mice infected with the Wuhan variant when the series of injections were started 12-24 hours post infection, and 50% survival for mice infected with the Gamma (P.1) variant when ACE2-V2.4-Ig was started 12 hours post infection [35]. In contrast, our data indicate that mice can be protected to a similar extent against the Delta variant using a 20-30-fold lower dose of ACE2-V2.4-Ig, and only one or two injections.

Several studies have also examined the ability of the LCB1 and LCB3 peptides, and derivatives, to protect mice from SARS-CoV-2 challenges. Using LCB1-Fc, which is equivalent to our SBP1-Ig, 100% of ACE2 transgenic mice survived challenge by the Wuhan virus, if 250μg of LCB1-Fc was given as IP injections 1 day pre- or 1 day post-exposure [39]. These doses are 25-fold higher than the dose we used here to study the effects of the tandem SBP1-1-Ig construct.

Our studies revealed the significant increase in potency and breadth achieved by linking 2 copies of the SBPs in the context of an Fc fusion, with SBP1-1-Ig being the most potent design we analyzed. Similarly, multimeric SBP constructs based on homotrimeric versions of another peptide, AHB2, proved to be highly effective in mice when given intranasally as a 50μg dose, 1 day pre-infection with either Alpha, Beta or Gamma variants, or 1 day post-infection with Beta or Delta [40]. Using this regimen, mice showed reduced weight loss and lower tissue viral loads by 6-7 days post infection, although the mice were not monitored long enough to look at effects on survival. In comparison, our SBP1-1-Ig construct protected 75% of animals from death over 11 days following only a single dose of 7.9 μg, given IP at 1 day post-infection with the Delta variant.

In summary, we have shown that ACE2-V2.4-Ig is a broad inhibitor of all current variants of SARS-CoV-2 and would likely be a good candidate as an alternative therapy to antibodies in the future as the virus continues to mutate. Multiplexed SBP constructs, such as SBP1-1-Ig, also hold promise as effective therapies since the multiplexed design overcame current circulating mutations that led to resistance against the monomers. However, the pattern of evolution of SARS-CoV-2 warrants that all such therapeutics will need to be monitored for effectiveness as new viral variants arise.

## Acknowledgements

This work was supported by a grant to PMC from a gift by the W.M. Keck Foundation to support COVID-19 research at the Keck School of Medicine of USC. Work with SARS-CoV-2 was performed in the Biosafety Level 3 Facility within The Hastings Foundation and The Wright Foundation Laboratories at the Keck School of Medicine and supported by a grant from the COVID-19 Keck Research Fund.

## Material and methods

### Plasmids and cell lines

ACE2-Ig comprises the extracellular domain of human ACE2 (amino acids 1-740) which was amplified by PCR from human cDNA using primers forward (5’-CAGTGTGGTGGAATTCACCATGTCAAGCTCTTCCTGGCT-3’) and reverse (5’-CACAAGATTTGGGCTCGGAAACAGGGGGCTGGTTAG-3’) and cloned into plasmid pVAX-IGHG1ΔCH1 by In-Fusion cloning (Takara Bio, San Jose, CA). This plasmid contains an Fc domain comprising Kabat numbers 226-478 of human IgG1. The V2.4 mutations (T27Y, L79T and N330Y) [28] were introduced using site-directed mutagenesis, to create ACE2-v2.4-Ig. For SBP1-Ig and SBP2-Ig, gBlocks (Integrated DNA Technologies, Inc., Coralville, IA) comprising the sequences of LCB1 and LCB3, respectively [35], were similarly inserted into pVAX with a (G4S)_3_ linker separating the peptide and IgG1 Fc domain. For SBP-Ig tandem repeat constructs, additional copies of peptide sequences were inserted, separated by (G4S)_3_ linkers.

An expression plasmid for Spike protein from the SARS-CoV-2 Wuhan-Hu-1 isolate (GenBank accession no. MN908947.3) in a pcDNA3.3 backbone was provided by James Voss (The Scripps Research Institute). SARS-CoV-2 variant Spike proteins were synthesized by PCR mutagenesis (D614G) or synthesized as gBlocks and cloned into the same expression plasmid, with the following variants obtained from the CDC variants tracker (https://www.cdc.gov/coronavirus/2019-ncov/variants/variant-classifications.html): Alpha-B.1.1.7, Beta-B1.351, Gamma-P.1, Delta-B.1.617.2, Epsilon-Cal20.C, Omicron-B. 1.537 or Omicron-BA.5.

293T cells and Vero E6 cells were obtained from ATCC. HeLa-ACE2 cells [50], were generously provided by James Voss. All cells were maintained in Dulbecco’s modified Eagle medium (DMEM) containing 10% fetal bovine serum (FBS; Gemini Bio, West Sacramento, CA) and 1% penicillin/streptomycin (Corning Inc., Corning NY), with Vero E6-ACE2 cells additionally supplemented with 1□μg/ml puromycin.

### Virus stocks

SARS-CoV-2 stocks were obtained from BEI Resources, NIAID, NIH: Wuhan isolate USA-WA1/2020, NR-52281, Beta isolate (B.1.351) South Africa/KRISP-EC-K005321/2020, NR-54008 and Delta isolate (B.1.617.2) USA/PHC658/2021, NR-55611. Virus was propagated on Vero E6-ACE2 cells as described [51] and stored at −80 °C.

### Production of entry inhibitors

Entry inhibitors were generated by calcium phosphate transient transfection of 293T cells, using 7μg/well plasmid DNA in 6-well plates. After 4 hours, media was changed, and supernatants collected after a further 48-hour incubation at 37^0^C. Inhibitor concentrations were determined by IgG ELISA. Briefly, goat anti-human IgG-Fc (Southern Biotech, Birmingham, AL) was coated onto 96-well plates overnight at 4^0^C. Plates were washed 3 times with PBS plus 0.05% Tween-20 (PBST) then blocked with 5% milk for 4 hours at 4^0^C. Serial dilutions of inhibitor supernatants were added and incubated at 4^0^C overnight, with recombinant human IgG1 (Sigma, St. Louis, MO) used as a standard. Plates were washed 3 times with PBST, mouse-anti-human IgG Fc-HRP (Southern Biotech) was diluted 1:2000 in blocking buffer and added and plates were incubated for 2 hours at 37^0^C. Plates were washed 3 times with PBST and SigmaFAST OPD (Sigma) added. Plates were read with a Mithras LB940 plate reader (Berthold Technologies, Bad Wilbad, Germany) to calculate sample concentration.

### SARS-CoV-2 Spike VSV pseudoviruses and neutralization assays

Replication-deficient VSVΔG vectors (76), containing an expression cassette for firefly luciferase and pseudotyped with various SARS-CoV-2 Spike proteins, were generated by transfection of 293T cells, followed by concentrated by ultracentrifugation or tangential flow filtration, as previously described [45]. The Spike pseudovirus stocks were titered by serial dilution on HeLA-ACE2 cells, with luciferase activity measured on a Mithras LB940 plate reader, as described [45], and calculated as relative light units (RLU).

For neutralization assays, 1 × 10^4^ HeLa-ACE2 cells/well were plated in 96-well half-area white plates (Corning, Inc., Corning, NY) and cultured overnight at 37^0^C. Spike pseudovirus stocks, calculated to produce approximately 100,000 RLU, were incubated alone or with different concentrations of inhibitors for 1 hour at 37^0^C before being added to the cells, in triplicate. The following day, media was removed and replaced with 50μl DMEM without phenol red and 25μl of Britelite plus (Perkin Elmer, Richmond, California), and luciferase activity was measured following the manufacturer’s protocol. Plates were read and RLU calculated using a Mithras LB940 plate reader. Best fit variable slope curves and IC_50_ calculations were calculated using PRISM software.

### SARS-CoV-2 plaque assays and addition of inhibitors

Plaque reduction assays were performed as described [51], and carried out in The Hastings Foundation and The Wright Foundation Laboratories BSL3 facility at USC. Briefly, confluent monolayers of Vero E6 cells in 6-well plates were infected with 100 PFU/well of virus and incubated at 37□°C for 45□min to allow virus adsorption, followed by replacement of the virus inoculum with media plus 1% low-melting agarose. For experiments in the presence of the ACE2-V2.4-Ig or SBP2-Ig inhibitors, virus was pre-incubated with the inhibitors for 1 hour at room temperature before addition to the cells. Plates were incubated at 37□°C for 72□hr, and plaques visualized by removal of agarose layer followed by 4% formaldehyde fixation and staining with 0.2% (w/v) crystal violet solution. Plaques were counted to determine plaque forming units (PFU).

### Mouse infections and treatment with inhibitors

Heterozygous K18-hACE2 C57BL/6J mice [47] were purchased from Jackson Laboratory (Bar Harbor, ME; cat. no. 034860) and transferred and housed within ABSL3 containment upon receipt. Female mice 9-10 weeks old were challenged through intranasal inoculation with 1×10^3^ PFU SARS-CoV-2 Delta variant (clinical isolate USA/PHC658/2021, NR-55611) in 30 μl PBS, by intranasal injection. Control mice received PBS alone. Mice were additionally treated with inhibitors at indicated times and doses, by intraperitoneal injection. Mice were monitored and weighed daily and euthanized due to declining health, including but not limited to 20% weight loss or failure of objective reflex tests. All animal care and experiments were performed according to the NIH guidelines for the care and use of laboratory animals and were approved by the Institutional Animal Care and Use Committee of the University of Southern California.

