## Supplemental figures 1-7 for "Comparison of SARS-CoV-2 entry inhibitors based on ACE2 receptor or engineered Spike-binding peptides"

**Running title:** SARS-CoV-2 entry inhibitors

*Address correspondence to:

Paula M. Cannon,

**The PDF file includes:** Figures S1 to S7

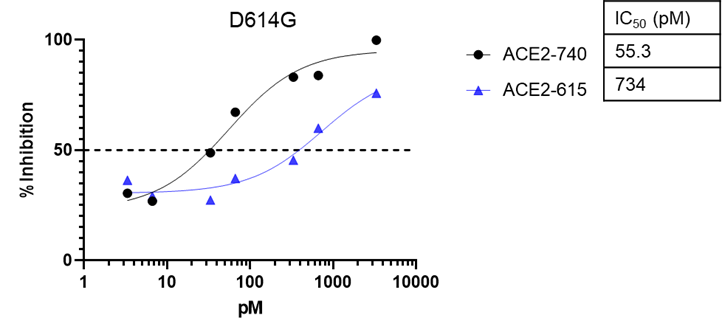

**Supplemental Figure 1. SARS-CoV-2 Spike neutralization by ACE2-Ig constructs with different ACE2 domains.** ACE2-Ig with ACE2 domains 1-615 or 1-740 were tested against VSV vectors pseudotyped with the D614G Spike protein and mean IC_50_ values are indicated.

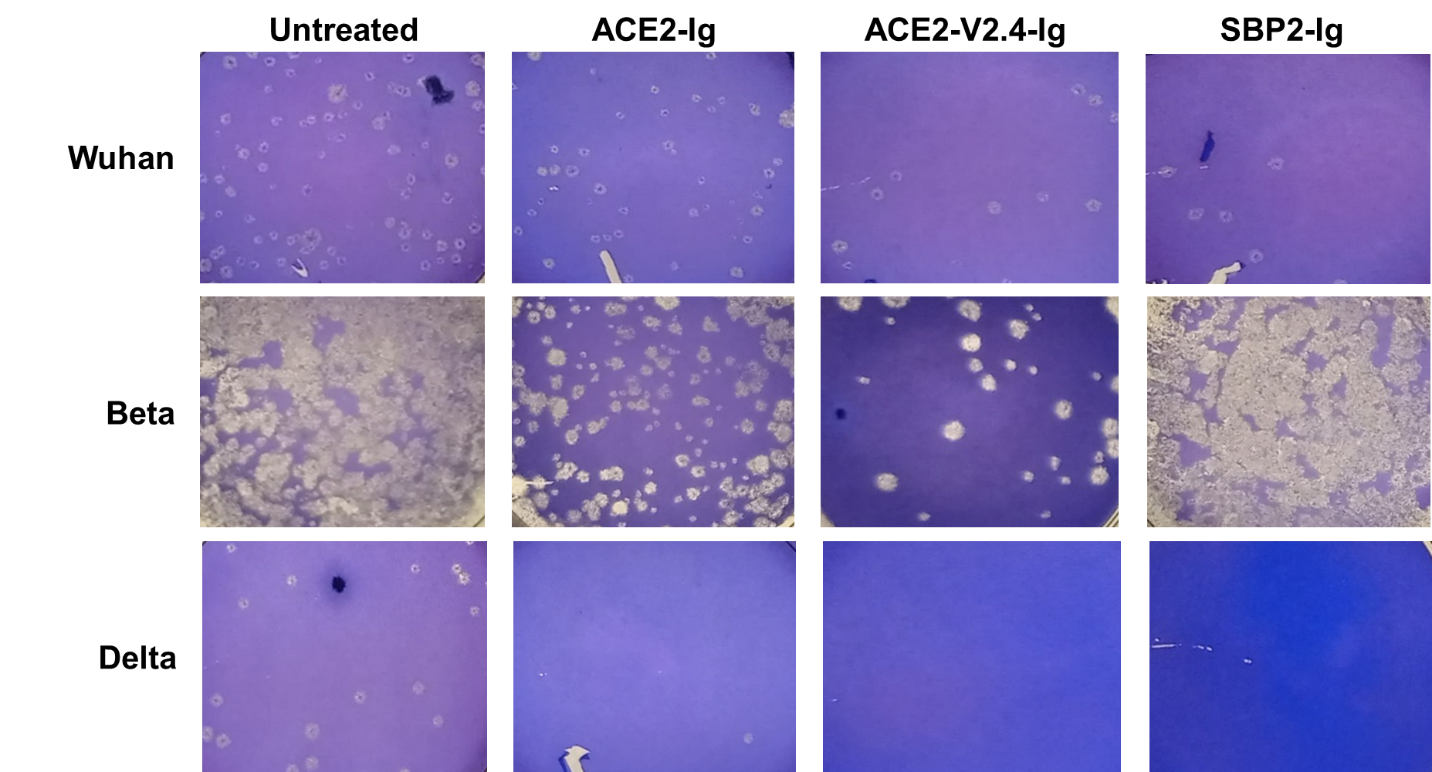

**Supplemental Figure 2. Activity of entry inhibitors against SARS-CoV-2 variants.** Qualitative plaque assays were used to assess the ability of ACE2-Ig, ACE2-V2.4-Ig and SBP2-Ig to inhibit the Wuhan, Beta and Delta variants of SARS-CoV-2.

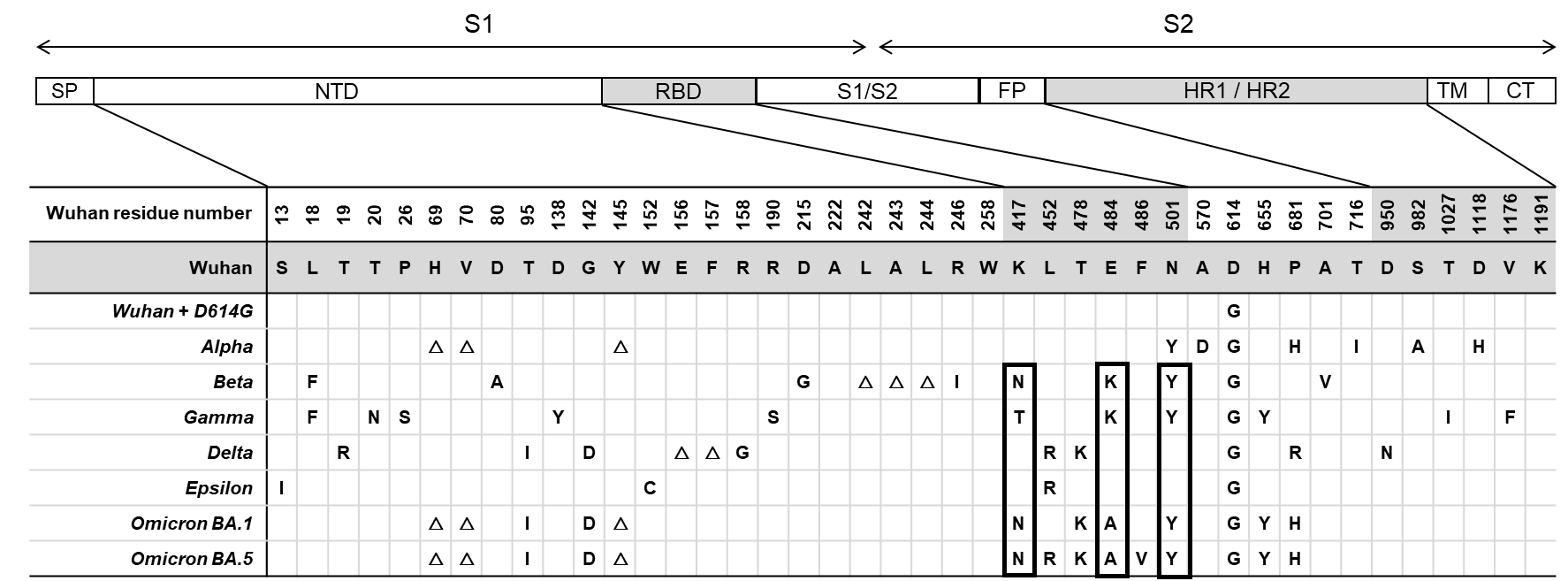

**Supplemental Figure 3**. **Mutations in the Spike proteins of SARS-CoV-2 variants.** Location of mutations in major SARS-CoV-2 variants in the indicated Spike protein domains and compared to the original Wuhan strain. Mutated residues in the RBD that are characteristic of the Beta, Gamma and Omicron variants are boxed. *Not shown, the Omicron variants also contain the following mutations: A67V, △ 143,145, △211, L212I, Ins214EPE, G339D, S371L, S373P, S375F, N440K, G446S, S477N, Q493R (except in BA.5) G496S, Q498R, Y505H, T547K, N679K, N764K, D796Y, N856K, Q954H, N969K, L981F. Abbreviations: SP, signal peptide; NTD, N-terminal domain; RBD, receptor binding domain; FP, fusion peptide; HR1/HR2, heptad repeat 1 and heptad repeat 2; TM, transmembrane; CT, cytoplasmic tail.

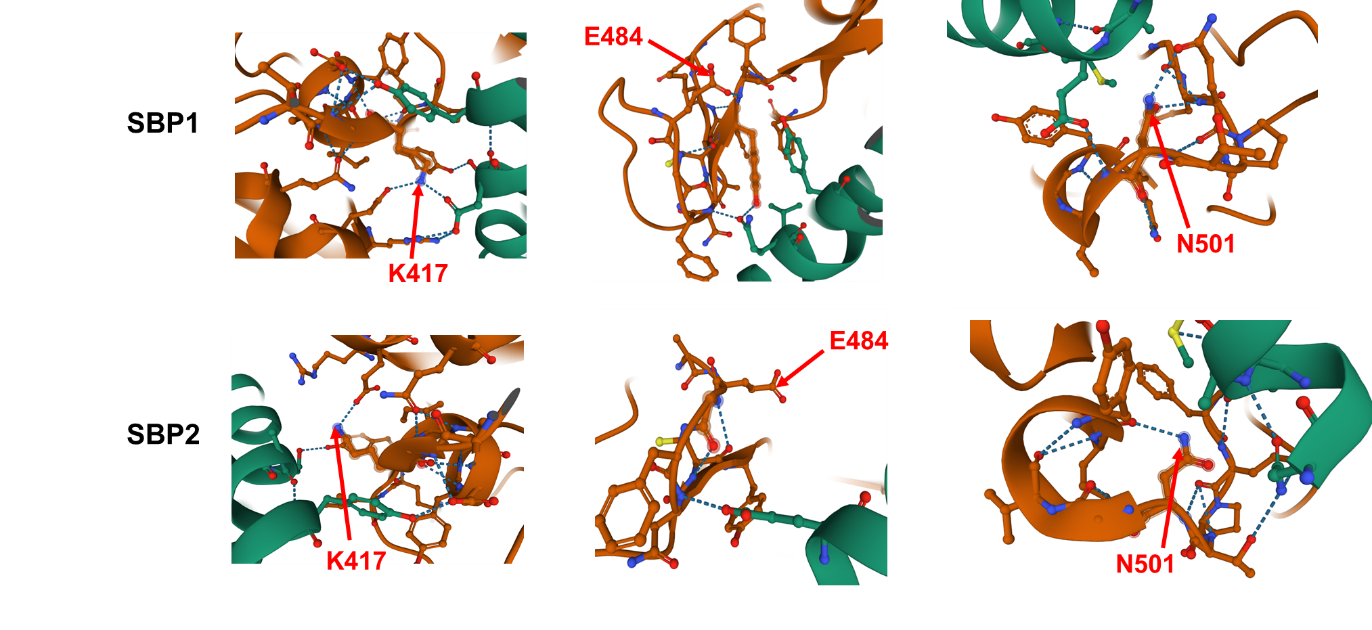

**Supplemental Figure 4.** SBP binding pocket in the RBD of SARS-CoV-2 Spike (Wuhan), showing relationship between Spike residues (green, with residues K417, E484 and N501 highlighted in red) and the two Spike-binding peptides (brown). Images generated from Protein Data Bank (PDB) website (<https://www.rcsb.org/>) for LCB1 (SBP1) in complex with SARS-CoV2 spike (PDB number 7JZU) and LCB3 (SBP2) in complex with SARS-CoV2 RBD (PDB number 7JZM.

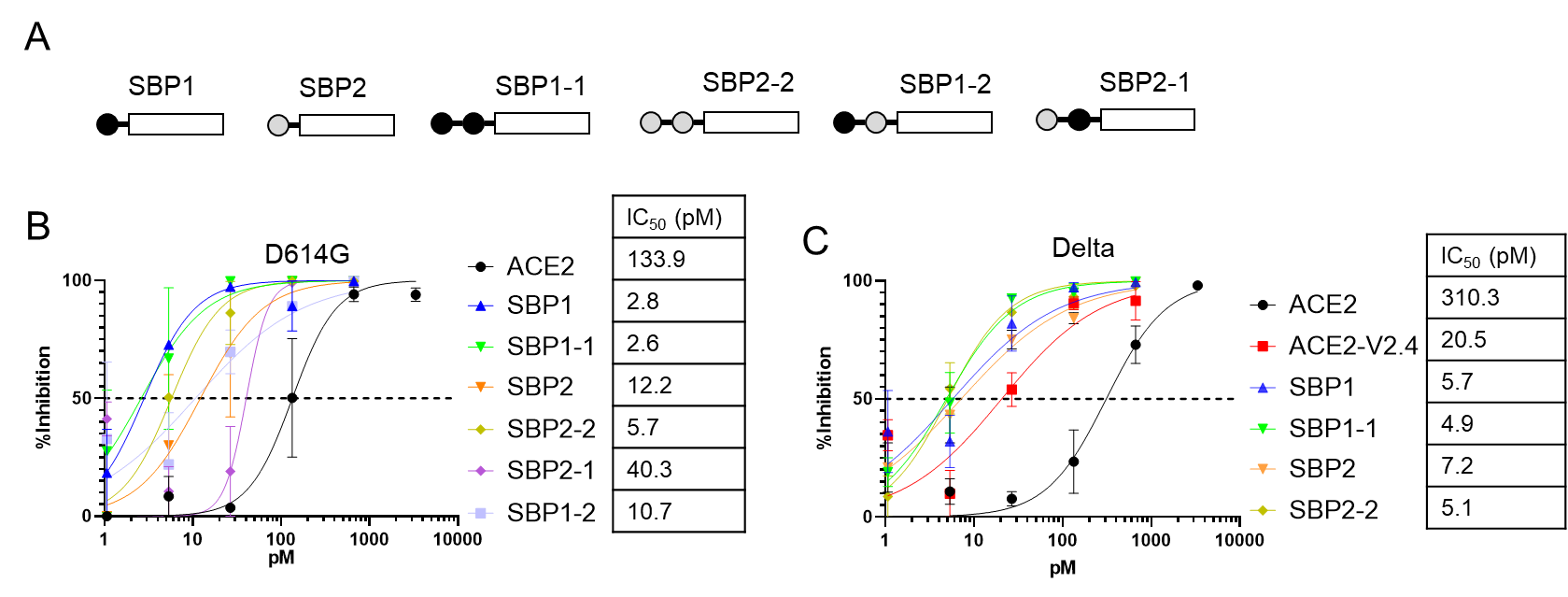

**Supplemental Figure 5. Effects of tandem combinations of SBP1 and SBP2 on D614G and Delta Spike neutralization. A)** Schematic of various SBP-Ig constructs. **B)** Neutralization assay against D614G Spike pseudovirus with indicated inhibitors. **C)** Neutralization assay against Delta Spike pseudovirus with indicated inhibitors. Error bars are SEM from n=3 independent experiments and mean IC_50_ values are indicated

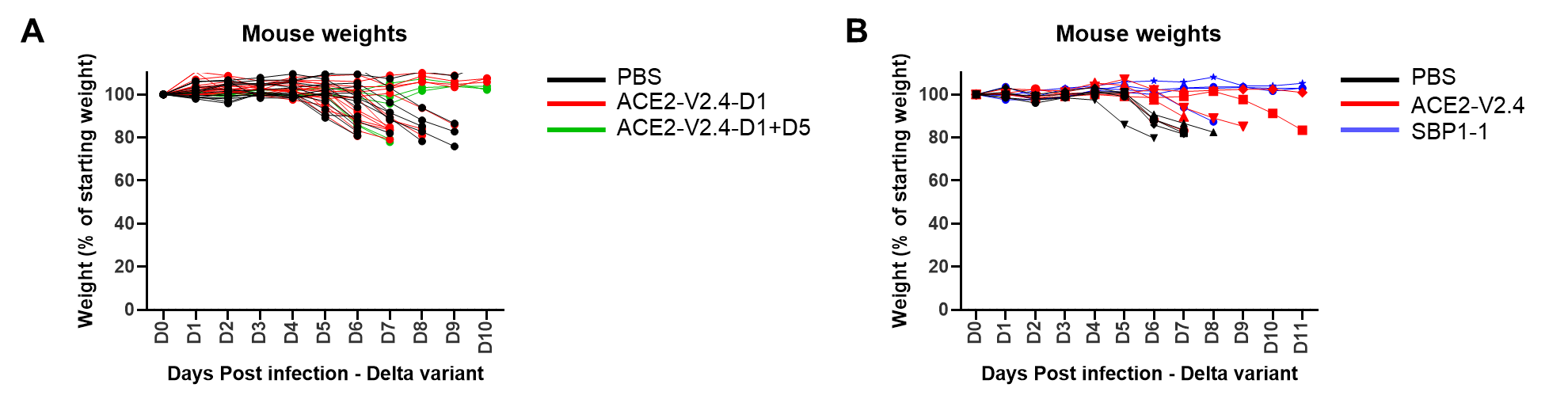

**Supplementary figure 6. Mouse weights over time during Delta virus infection.** Individual mice used in Figure 6 are plotted. **A)** Mice used in Figure 6A. **B)** Mice used in Figure 6B.

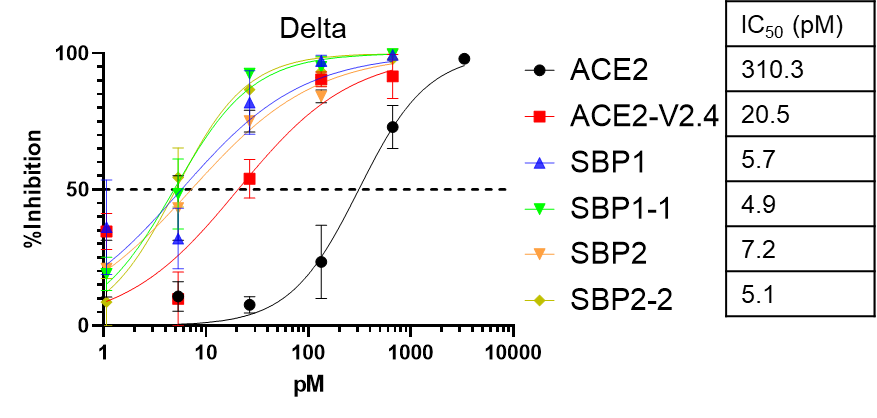

**Supplemental Figure 7. Tandem repeat SBP constructs inhibit Delta variant but not significantly more than SBP monomer constructs.** Neutralization assays were performed against the SARS-CoV-2 pseudovirus Delta variant using the indicated inhibitors and mean IC_50_ values are indicated. Error bars are SEM from n=3 independent experiments.
